# Phylogeny and gene function integration uncovers multiple convergences in multicellular and terrestrial transitions

**DOI:** 10.1101/2025.09.19.677313

**Authors:** Gaurav D. Diwan, Paschalis Natsidis, Mu-en Chung, Juan-Carlos Gonzalez Sanchez, Roderic Guigó, John K. Colbourne, Maximilian J. Telford, Robert B. Russell

**Author notes:** Corresponding authors: Gaurav D. Diwan Robert B. Russell.

## Abstract

In the biodiversity genomics era, vast gene catalogs have rapidly accumulated, but how gene functions map onto major evolutionary transitions remains vague. We integrated phylogenomics with functional annotations to trace the emergence of ∼4.5 million genes and their associated functions in 508 representative species across the tree of life. Using this top-down, function-centric approach, we mapped the evolutionary history of well-annotated biological functions, revealing key insights and widespread parallelism. For instance, the largest bursts of gene and function gains aligned with key evolutionary transitions, including the origins of eukaryotes, animals, land plants, and vertebrates. Independent transitions to multicellularity and terrestrial life showed convergent enrichment of semantically similar functions (involving entirely different gene families), and the presence of orthologous genes often belonging to ancient gene families. For example, terrestrialization nodes in slime molds, plants, and animals coincided with gains in stress-tolerance and developmental genes, whereas multicellularity in animals, fungi, plants, and slime molds consistently involved expansions in cell-adhesion and communication functions. Thus, our framework enables scalable interpretation of new genomes in an evolutionary context and offers a roadmap for exploring the genomic basis of biodiversity.

## Introduction

We are now in the biodiversity genomics era, marked by massive international projects cataloguing life’s genetic diversity at unprecedented scale^1–3^. Building on landmark efforts like the Human^4^ and Mouse^5^ Genome Projects, these initiatives have driven major advances in genome assembly, annotation, and comparative analysis^6,7^. While whole-genome sequencing and comparative analyses are now routine, major challenges remain in this era^8^, especially in assigning and interpreting gene function in an evolutionary context.

To use these expanding genomic resources fully, we must move beyond cataloguing genes to understanding what they reveal about the biology and evolution of organisms. How can we use the depth and breadth of sequenced genomes to reconstruct the deep past of life on Earth? And how can we use that perspective to identify which genes and gene functions are linked to particular phenotypic (clade-specific) traits, and when in evolutionary history they arose? Answers to such questions would deepen our understanding of major evolutionary transitions^9^, such as the emergence of eukaryotes, transitions to new environments, multicellularity, and cell type diversity, and illuminate core biological processes like development, regeneration, and even human-specific conditions including cancer and aging. Realizing the potential contained within the data we now have requires a robust framework for tracing functional annotations across the tree of life.

Phylogenomic analyses provide an avenue for this by enabling inference of the gene content and phenotypic traits of ancestral species^10–12^. Previous studies have successfully reconstructed ancestral genomic content either for a handful of genes/functions or for specific clades^13–23^, and specialized databases such as OMA^24^, EggNOG^25^, STRING^26^, PhylomeDB^27^, and Matreex^28^ provide phylogenetic-profile–based views of orthology and function. While some of these efforts have traced the evolutionary history of all genes within a given clade, they typically lack the broader phylogenetic context needed to distinguish between truly clade-specific innovations and older, conserved families that pre-date the clade itself. Without considering sister lineages, it remains unclear whether genes gained at the apparent root of a clade represent genuine novelties or inherited ancestral families. Likewise, studies focusing on particular pathways or gene sets illuminate the evolution of those specific functions but overlook other functions gained at the same nodes, information essential for identifying potentially co-evolving or functionally coupled innovations. These gaps underscore the need for a large-scale framework that reconstructs the evolutionary history of all genes and functions simultaneously across the entire tree of life. Such a framework can then reveal whether functions are restricted to certain clades, repeatedly reinvented in independent lineages, or useful as guides for annotating newly sequenced species.

To explore these opportunities, we asked whether combining phylogenetic relationships with functional annotations across the tree of life could identify the evolutionary emergence of widely understood traits and functions. Moreover, whether we could use the evolutionary history of genes and their functions to inform us of the genomic events that coincided with large phenotypic transitions. In this study, we analyzed genomic annotations as available in public databases, in the context of a deeply rooted phylogeny of life comprising 508 representative species (Table S1; Materials and Methods). Combining a custom-built species tree with OrthoFinder^29^, we sorted ∼4.5 million proteins into hierarchical orthologous groups (HOGs) defined at the root; and traced the evolutionary history of each gene (using the presence/absence of their HOGs as well as individual orthologs) and its associated functions using Gene Ontology terms^30,31^, protein domains (Pfam^32^), and biological pathways (KEGG^33^; Reactome^34,35^).

### The largest gain events correspond to specific biological traits

Of the largest gain events we detected (Fig. 1, Table S2), most were confined to the eukaryotic superkingdom: in the ancestors of eukaryotes, slime molds (Evosea), land plants (Embryophyta and Magnoliopsida), animals (Metazoa) and vertebrates (Chordata and Euteleostomi). Among the prokaryotes, two large gain events are apparent in Cyanobacteria and one in Archaea (Halobacteria). For most founder events, genes (i.e. HOGs; see Materials and Methods), protein domains and pathways were gained in large quantities simultaneously (Fig. S2). In contrast, gene losses occurred six times less frequently than gains (Fig. S3; Fig. S4A), indicating that expansion, rather than reduction, dominates the large-scale evolutionary signal.

**Fig. 1:**
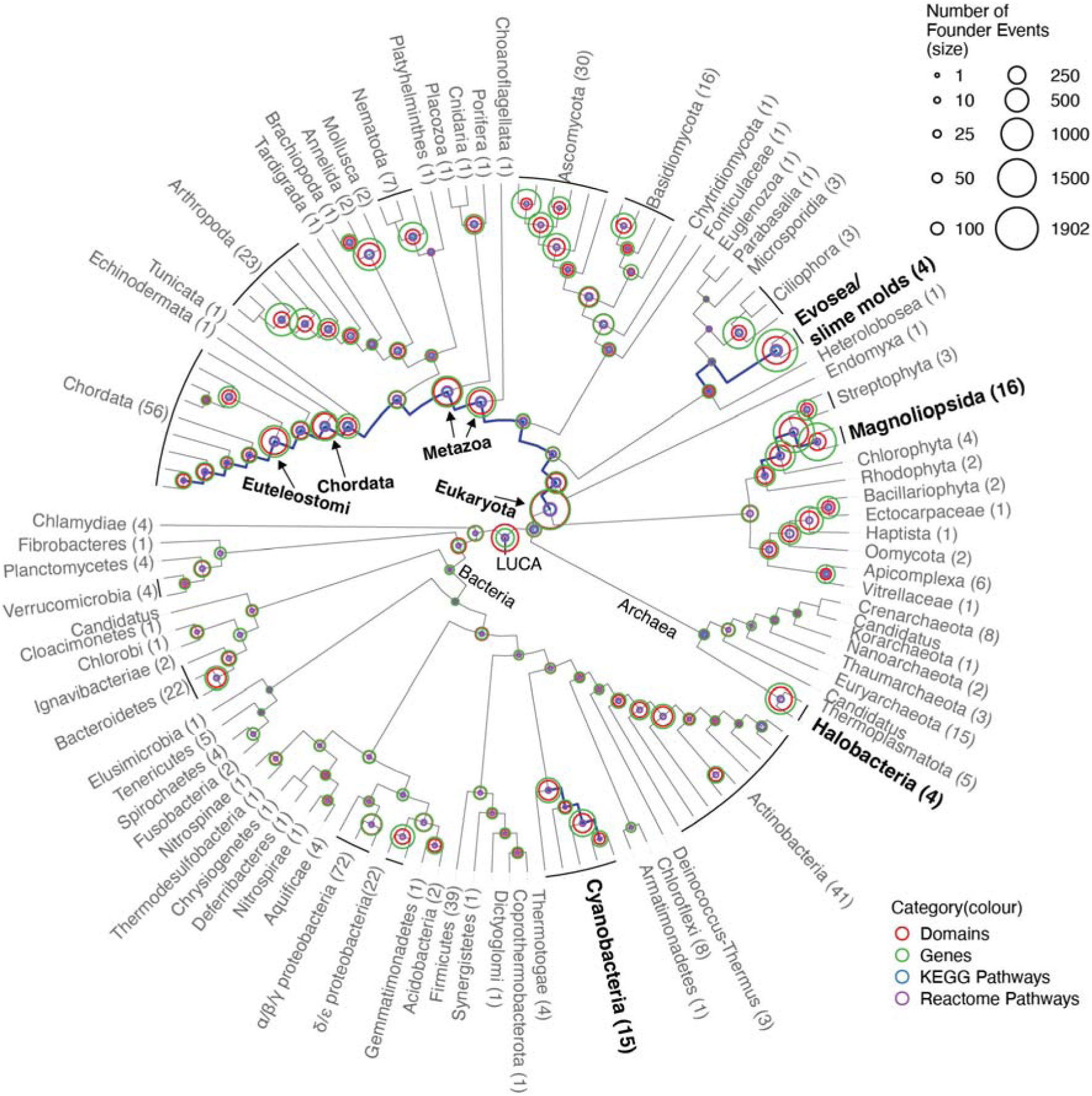
Ancestral reconstruction of genes, domains and pathways reveal nodes with the largest gain events. Phylogenetic tree (cladogram) showing the founder events of biological entities in the ancestors of the clades in this study (number of species in parentheses; the complete tree with support values is shown in Fig. S1). The size of the coloured circles represents the number of each category that was gained at a given node. N.B.

KEGG/Reactome pathways are depicted as the sum of the proportion of pathways gained at the node. Lineages with large and simultaneous emergence of multiple biological categories have been emphasized (dark blue lines). These are further detailed in Fig. 2. LUCA = Last Universal Common Ancestor

**Fig. 2:**
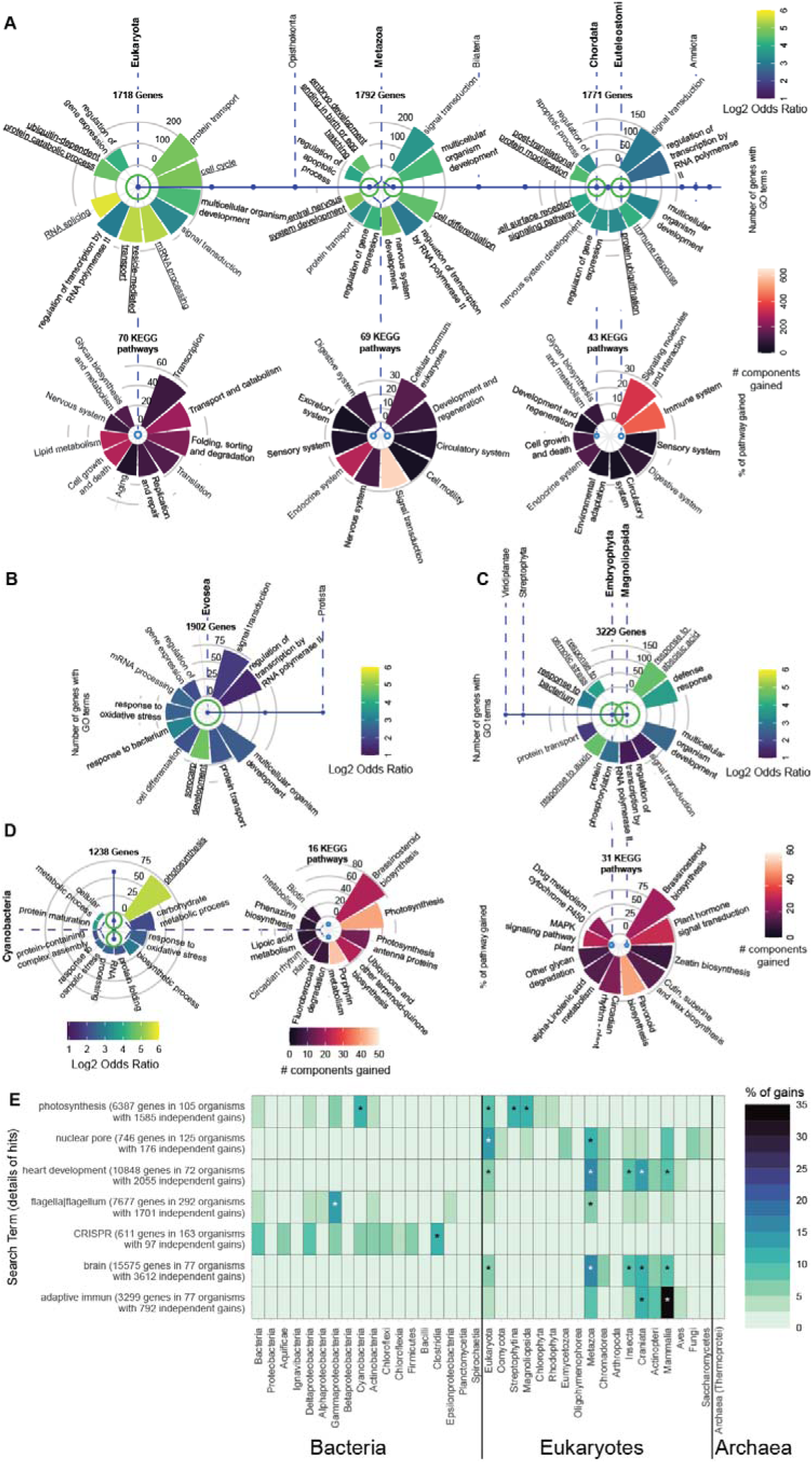
Combining phylogeny with functional and pathway annotations shows gains of well-known traits at different nodes in the tree. A-D) Figures showing a detailed look at the largest founder events from Fig. 1. The highlighted lineages (dark blue lines) in Fig. 1 have been depicted in a linear manner and have been labelled with the relevant taxonomic information wherever possible (dashed lines). In each panel, bar plots show the ten enriched GO terms with the highest number of genes in each category and the ten KEGG pathways with the highest % of the pathway gained at the respective node in the tree. The GO term bars are coloured by the Log2 Odds Ratio from a Fisher’s exact test and the KEGG pathway bars are coloured by the number of components (i.e., proteins) gained for each pathway. Unique GO terms from each clade are also underlined. E) Heatmap showing the % of gains for the genes captured by a search term occurring at each of the nodes on the x-axis. Asterisks indicate the taxa that have significantly higher % gains than the mean (z-score > 1.96) calculated for each search term independently.

Linking gene gains to function revealed patterns that closely matched the emergence of major biological traits. To quantify functional enrichments, we adapted standard GO enrichment analyses by counting the number of HOGs containing genes annotated with each term, rather than the total number of genes. This approach better reflects independent gene-family contributions to a function and reduces biases from large gene families. For pathway analyses, we focused on the broader levels of the KEGG hierarchy (H1 and H2), which provided more informative summaries across taxa. Using these approaches, we found that the eukaryotic root gained a large proportion of fundamental pathways such as transcription (on average 54.6% of KEGG pathways in this category) and translation (37.4%), and showed enrichment of broad GO terms including cell cycle, mRNA processing, and RNA splicing (Fig. 2A). The metazoan root gained pathways including development and regeneration (30.9%), circulatory system (30.1%) and cell motility (29.1%), and was enriched in terms such as cell differentiation and embryo development (Fig. 2A). Further specificity of function was observed in the vertebrate ancestor with large gains in pathways for signalling processes (27.7%) and the immune (24.6%), sensory (18.7%) and digestive (18.7%) systems, and in terms such as ‘immune response’ and signaling pathways (Fig. 2A). The ancestor of angiosperms showed an enrichment of GO terms such as ‘response to abscisic acid’, ‘auxins’ and ‘bacteria’ and pathways such as Brassinosteroid synthesis (92.8%), plant hormone signal transduction (79.5%), plant circadian rhythm (70.5%) and zeatin biosynthesis (69.2%) (Fig. 2C) whereas the slime molds showed an enrichment for sorocarp development (Fig. 2B).

At the same nodes that showed the largest bursts of gene gains, we also examined the corresponding gene losses (Fig. S4B). These involved far fewer HOGs (tens versus hundreds for gains) and had narrower functional scope. Losses at Metazoa and Magnoliopsida were enriched for translation, mitochondrial translation, RNA processing, and lipid metabolism, suggesting minor streamlining of core processes. In slime molds, enrichment of gene losses included ciliary and developmental terms (e.g., cilium assembly, gastrulation) reflecting for example the known loss of cilia in this clade^36^, while in Arthropoda and Craniata, losses affected diverse functions such as DNA repair, respiration, and signaling. Overall, these particular nodes were dominated by functional innovation rather than reduction.

By contrast, the nodes with the largest number of losses overall included parasitic or genome-reduced eukaryotes such as Apicomplexa, Nematoda, Microsporidia, and diverse unicellular taxa (*Trichomonas*, *Leishmania*, *Entamoeba*, several ciliates), as well as mycobacteria (Fig. S3). Here, GO enrichment of lost genes showed convergent patterns of simplification. Most nodes lost genes involved in signal transduction, multicellular development, protein transport, transcription regulation, and mRNA processing, consistent with genome streamlining. Mycobacteriales and Lactobacillales lost respiratory and biosynthetic genes as well as transmembrane transport (Fig. S4C). Thus, while losses trace specialization and simplification, gains capture broad episodes of functional innovation.

To generalize our approach beyond curated functional sets, we used keyword-based text queries across annotations (Fig. 2E). For each term we identified clades where genes containing the term of interest in their annotations originated. For instance, the genes containing the term “photosynthesis” in their GO annotations originate mostly in eukaryotes and plants (Viridiplantae and Embryophyta, Fig. 2E). A few genes originated in bacteria, with the largest gains in cyanobacteria (Cyanophyceae). Terms such as “nuclear pore,” “heart development,” “brain,” and “adaptive immun(e)” showed genes with gains in eukaryotes and their descendant clades, often peaking in vertebrates (e.g., adaptive immunity genes in Euteleostomi and Primates). Genes annotated with “flagella|flagellum” originated independently in bacteria and eukaryotes, reflecting their presence in bacterial motility genes and ciliary structures such as intraflagellar transport complexes. As expected, genes associated with “CRISPR” were confined to bacteria and archaea.

Together, these findings demonstrate how combining phylogenomic inference with large-scale functional annotation can effectively recapitulate the biology of diverse lineages. The enrichment of pathways and GO terms at key ancestral nodes mirrors the defining innovations of those clades, showing that annotation data alone, when integrated with phylogenetic context, can illuminate the emergence of complex traits. Having established that this framework reliably captures known evolutionary signatures across the tree of life, we next apply it to investigate whether similar patterns of coordinated gene and function gains accompany specific phenotypic transitions, such as the move to land and the origin of multicellularity.

### Semantically similar functional terms are repeatedly enriched in multicellularity and terrestrial transitions

To establish a broad view of the genomic changes accompanying major phenotypic transitions, we asked: when lineages made the shift to land or to multicellular life, did their newly gained genes converge on similar functional categories? To test this, we applied the recently described framework of phylogenetic genotype-to-phenotype mapping^37^. We identified phylogenetic nodes that underwent the respective transitions (Fig. 3) and enriched GO terms among HOGs gained at each transition node. Next, we clustered these terms by semantic similarity (using Lin similarity^38^), and compared their enrichment to that of their parent nodes (see Materials and Methods; Fig. 4A). Each GO cluster represents a set of functionally related terms linked through shared parent terms in the GO hierarchy. Several clusters revealed biologically coherent patterns that, in retrospect, match known adaptive pressures associated with these transitions. An equivalent exercise using pathway information proved uninformative largely because of limited (or even entirely flat) hierarchies, in contrast to the deep hierarchical structure of GO.

**Fig. 3:**
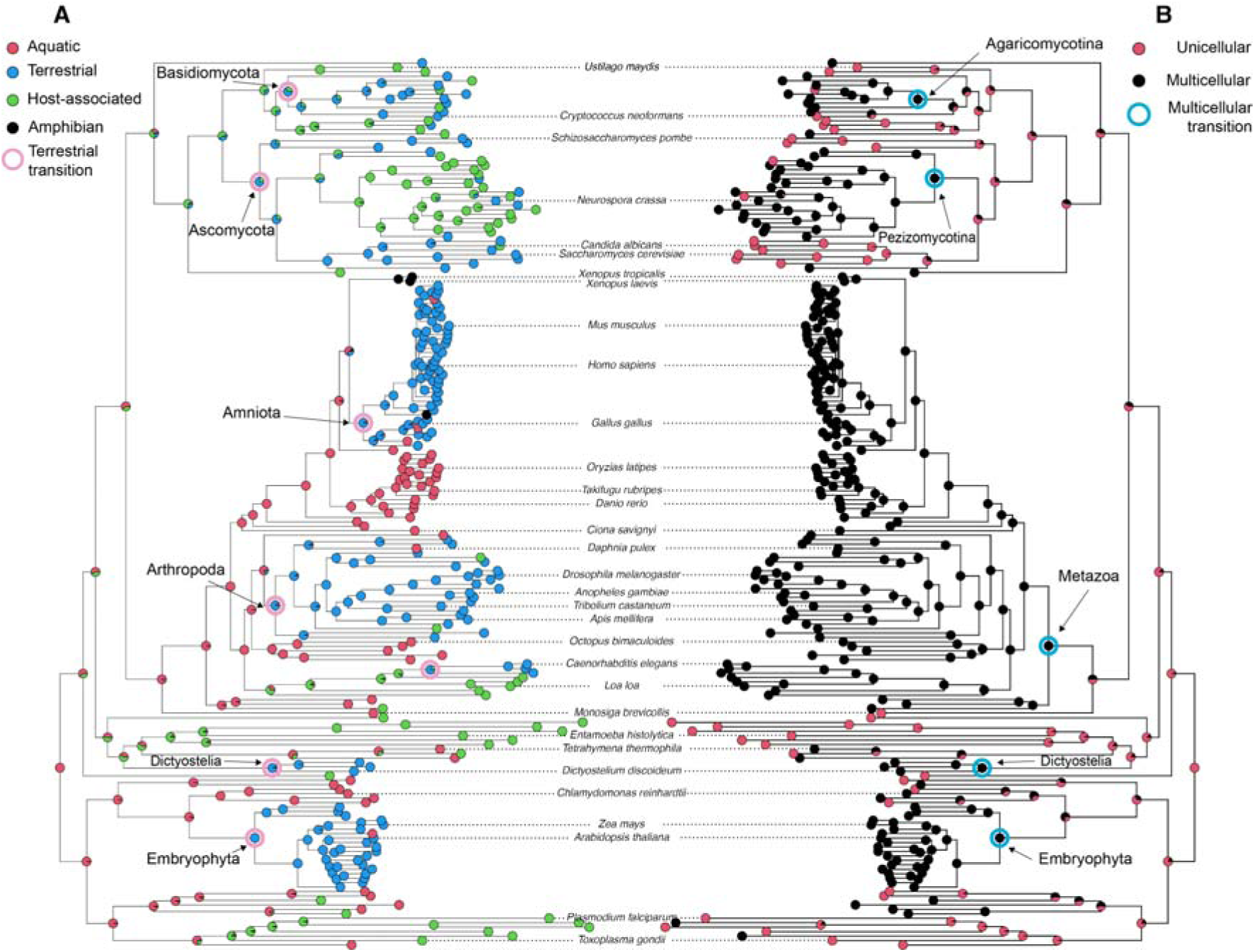
Ancestral reconstruction reveals nodes that underwent transitions to land and multicellularity. Stochastic character mapping of broad phenotypic traits encoded for the tips of the eukaryotic sub-tree from Figure 1. Each node depicts a pie chart of the posterior probabilities for each character state. Nodes that went from predominantly aquatic or host-associated/unicellular states to predominantly terrestrial/multicellular states have been highlighted (posterior probability cutoff of 0.75 and 0.5 respectively). A subset of well-studied species have been labelled for orientation. N.B. The Host-associated state includes pathogens and commensals; and the Terrestrial state includes airborne species.

**Fig. 4:**
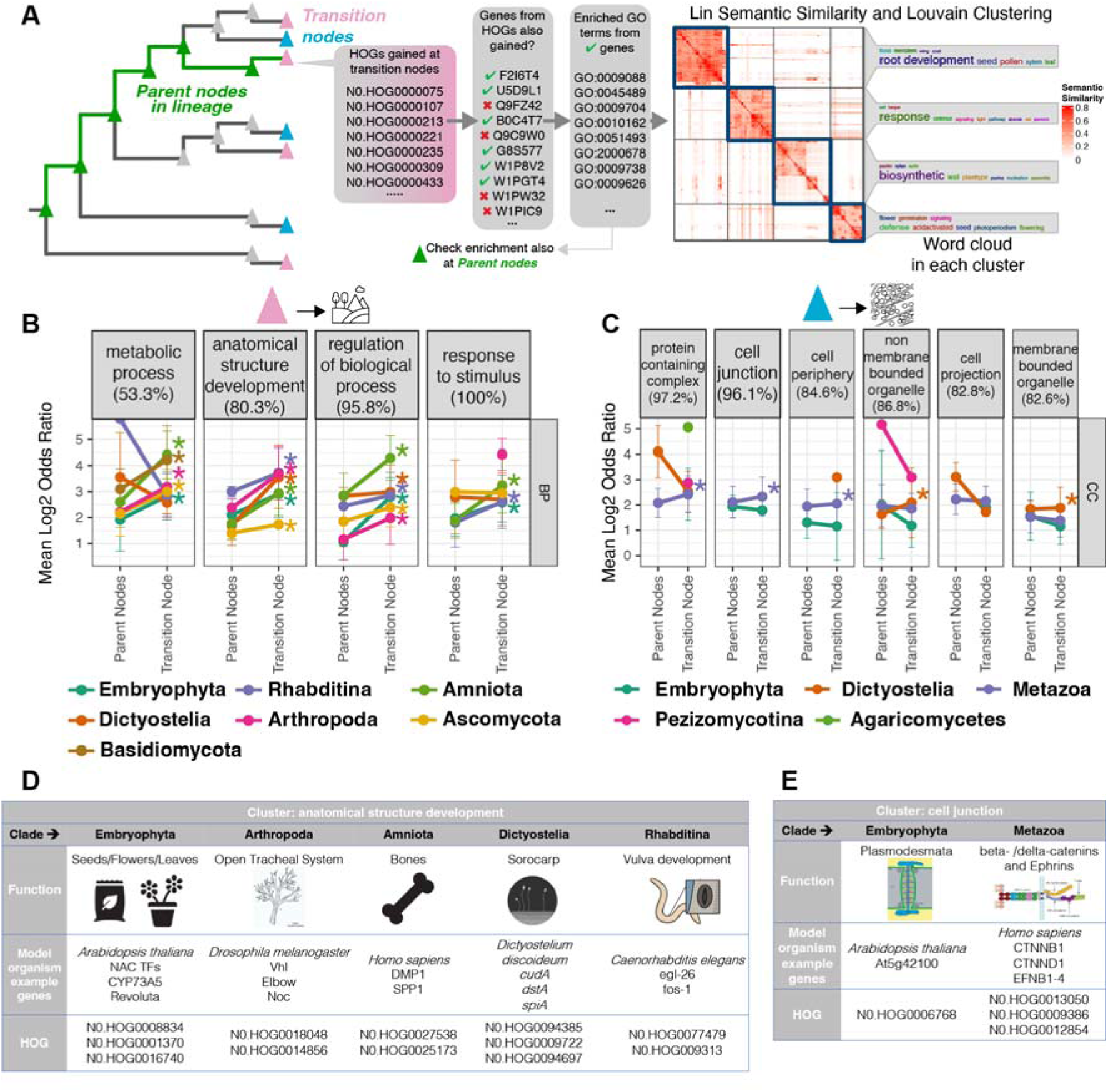
Functional clusters that include adaptive genes and traits for large phenotypic transitions are enriched at transition nodes. A) Schematic representation of the GO term analysis done to elucidate the general functions gained at each transition node. We used the HOGs gained at each node and took the GO terms of the gene members from these HOGs that were also gained at the same node. We carried out an enrichment analysis of these terms at the transition nodes as well as the parent nodes for each lineage. The enriched terms were then clustered using Lin Semantic Similarity and Louvain Clustering and we manually determined the most informative common parent GO term for labelling the clusters. B-C) Mean (and Standard deviation) of Log2 Odds Ratio for the significant GO terms in the different clusters at transition nodes compared with parent nodes. The most informative parent terms encompassing the GO terms in each cluster were used to label the panels (% of terms in the cluster with the respective parent term is shown in parentheses). Asterisks indicate a significantly greater mean of Log2 Odds Ratio at the transition node. We discuss the GO:BP (biological process) clusters for the terrestrial transition and GO:CC (cellular component) clusters for the multicellular transition; other categories are shown in Fig. S5. D-E) Tables showing genes from well-studied model organisms that are involved in the indicated function and gained at the terrestrial and multicellular transition nodes respectively. These genes were part of distinct HOGs indicating that they were clustered via their functional annotations and not sequence similarity.

For the terrestrial transition, we identified seven clades in our phylogeny that independently colonized land: Dictyostelia, Ascomycota, Basidiomycota, Embryophyta (land plants)^39^, and animal clades such as Rhabtidina (nematodes), Arthropoda^40^, and Amniota (tetrapods excluding amphibians)^40,41^ (Fig. 3A; Table S2). Other transitions such as in tardigrades, rotifers, molluscs and onychophorans^40^ were not captured due to limited sampling of these clades in our study. For the seven transition nodes we detected, enrichment was observed in functional categories related to environmental adaptation and structural innovation. One major GO:BP (biological process) cluster included terms associated with anatomical structure development (parent term of 80.3% GO terms in this cluster) and was significantly enriched across six transition nodes (all except Basidiomycota). This enrichment was also significantly greater than in the corresponding parent nodes (Fig. 4B; Table S3). Organisms moving to land evolved several convergent phenotypic adaptations such as the water retentive structures and skeletal changes^42^. Interestingly, genes in the anatomical structure development cluster encoded for such functions. For instance, in tetrapods, the cluster contained osteopontin (SPP1) and dentin matrix acidic phosphoprotein 1 (DMP1), both involved in bone development; in plants, we observed regulators like NAC transcription factors involved in seed, leaf and flower development (e.g., NAC034, NAC051, NAC098), lignification-related CYP73A5, and vascular patterning genes such as class III HD-Zip genes (e.g., Revoluta); in arthropods, gains included components of the open tracheal system such as Protein Vhl, Zinc Finger Protein Elbow, and Noc (Fig. 4D; Table S4). Going beyond these well studied clades in terrestrialization studies, we also observed genes from Dictyostelia such as *cudA*, *dstA* and *spiA* that are involved in sorocarp development (enclosed structures carrying spores); and the egl-26 and fos-1 genes in nematodes leading to the development of the vulva (a structure aiding in internal fertilization on land; Fig. 4D; Table S4). Importantly, most if not all of the genes (68125 out of 68310) in this cluster belonged to different HOGs (Fig. 4D; Fig. S6), showing that the convergence via their functional annotations pointed to common adaptive strategies rather than their sequence similarity.

Another major cluster linked to terrestrial transitions involved response to stimuli (all GO terms in this cluster had this parent term). This cluster was significantly enriched at all terrestrial transition nodes except Basidiomycota (Fig. 4B; Table S3). Organisms moving to land faced new environmental conditions, including exposure to pathogens, reduced water availability, along with a substantially altered light spectrum. Indeed, in plants, this cluster included genes involved in red/far-red light response such as *Empfindlicher im dunkelroten Licht* protein 1 (EID1), as well as abscisic acid signaling genes (e.g., ABI5, ABF1–4), which mediate responses to desiccation and other land-specific stresses. In land animals, this cluster encompassed expansions in components of the adaptive immune system^43^, including interferons and LY96, reflecting increased pathogen defense complexity. For nematodes, this included genes such as abf-2 which encodes an antimicrobial peptide and in arthropods, genes involved in circadian rhythm entrainment such as *per* and *tim* (Table S4).

We applied a similar approach to the emergence of multicellularity, identifying independent transitions in Metazoa, Dictyostelia (slime molds), Embryophyta, and multicellular fungi (Agaricomycotina and Pezizomycotina) (Fig. 3B; Table S5). In this transition, we expected functional enrichment related to cell–cell communication and coordination. Indeed, we observed a GO:CC (cellular component) cluster involving cell junction terms like “synapse,” “adherens,” and “junction” in both land plants and Metazoa (Fig. 4C). In Metazoa, this cluster included beta- and delta-catenins and Ephrins, which mediate intercellular adhesion and signaling (Fig. 4E; Fig. S7; Table S6). In Embryophyta, the same cluster included genes involved in plasmodesmata formation, the plant-specific channels for intercellular communication (Fig. 4E). The transition to multicellularity also required substantial increases in Golgi-mediated secretion and vesicle trafficking^44^, supporting the expanded extracellular matrix, adhesion proteins, and signaling molecules needed for coordinated cell–cell interactions. Interestingly, another GO cluster we detected was for membrane bounded organelles (Fig. 4E) and included terms related to the Golgi apparatus and secretory granules across land plants and animals (Table S5). This cluster included genes such as SNARE family protein At1g71940 from *Arabidopsis thaliana* and the secretory granule gene Snap24 from *Drosophila melanogaster* (Table S6).

In summary, the enrichment analyses reveal that broad categories of function viz. anatomical structures, developmental processes, stress responses, and cell–cell junctions, emerged repeatedly in lineages undergoing similar transitions, and point to common adaptive solutions even though the underlying genes differ. This suggests that convergence can be detected at the functional annotation level before focusing on specific molecular players. Indeed, we also next asked whether previously described “transition genes” follow the same pattern of emerging around the transitions, and when the gene families containing them first arose.

### The timing of gains for known genes across transition nodes reveals co-option of ancient gene families

Having established broad functional signatures, we next focused on specific candidate genes and their families that have been previously linked to these phenotypic transitions. Our approach had the advantage of timing the appearance of genes and their gene families (i.e. HOGs) along the phylogeny, allowing us to ask whether the origin of particular HOGs coincided with, or preceded, the evolutionary nodes corresponding to major phenotypic transitions. Specifically, we asked whether genes previously described as playing a role in these transitions within certain clades are also present as orthologs in other clades that underwent the same transitions independently, and when their broader gene families first emerged. To do this, we reconstructed a timeline of both the presence of orthologs for these known genes and the points at which the corresponding HOGs were gained, using the first inferred gain of a HOG as a proxy for the origin of that gene family.

For the terrestrial transitions, we focused on genes previously associated with adaptation to land (Table S7) including BMP4, Noggin, Pax2, Eyeless/Pax6, KEAP1/NRF2, Hox genes, and Aquaporins^41,45,46^. These genes have been greatly described previously and we chose them because they regulate body patterning, organogenesis, vision, oxidative stress response, and water balance, traits that were critical for adapting to terrestrial environments. In clades where these genes have previously been described viz. tetrapods, arthropods, and land plants, orthologs of genes were detected near or before the respective transition nodes, often derived from HOGs gained earlier in the tree (Fig. 5A; Fig. S7; Table S8). Significant enrichment points (asterisks; one-sided binomial test, FDR < 0.05) identified nodes where the fraction of known transition genes inferred as present exceeded the genomic expectation, indicating specific gains of these genes at transition relevant points of the phylogeny (Table S8). For instance, Hox genes and Aquaporins show punctuated enrichments around the tetrapod transition, while GRAS transcription factors rise similarly at the plant transition (Fig. 5A). Smaller gene sets such as Noggin, Pax2, or BMP4 show analogous qualitative peaks despite limited statistical power.

**Fig. 5:**
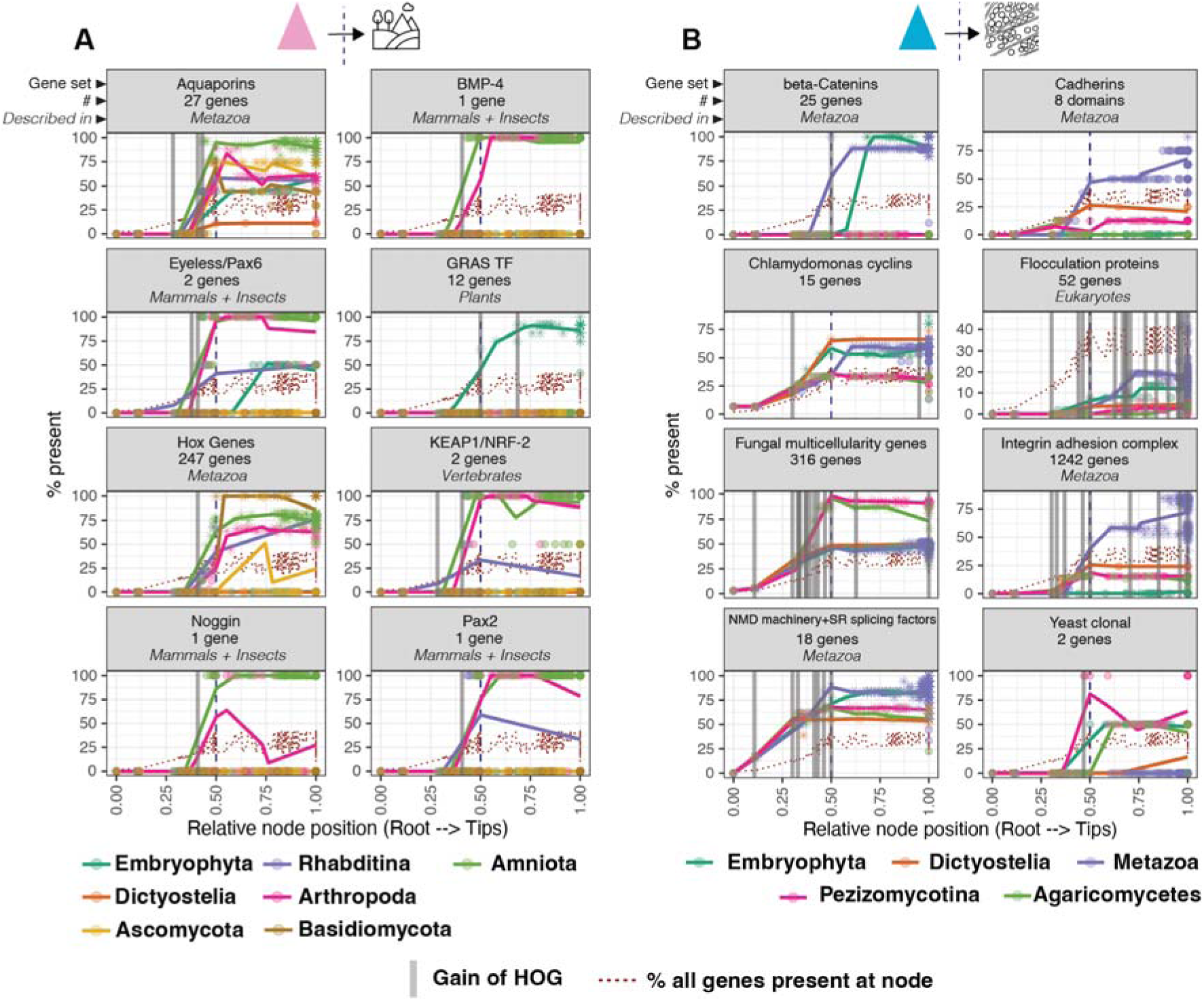
Known genes previously described for large phenotypic transitions are present parallely in multiple transition lineages. A-B) Plot showing the % of orthologs present (out of the number of genes/domains indicated for each set) at each node of a transition lineage. The clades where each gene set was previously described in the literature is indicated in italics. The dotted brown line indicates the genomic expectation at each node used in these analyses (% of all genes inferred to be present at a node). We tested if the % present at every node for each gene set was significantly greater than this genomic expectation using a one-sided binomial test and points that were significant are shaped like an asterisk. The distance to the root of each node in a lineage was scaled between 0 and 1 on the x-axis, while the transition nodes were forced to be at 0.5 (dashed vertical lines). To indicate the trend of the presence, multiple linear models were fitted to the underlying points for each lineage (one model each for four equal sections of the x-axis, see Materials and Methods). Vertical gray bars indicate the earliest gain of HOGs associated with the known genes.

Notably, orthologs of Eyeless and Pax6 were also detected in land plants^47^ and nematodes respectively, but from earlier-emerging HOGs, highlighting parallel use of ancient gene families across lineages. Metazoan aquaporins taken from a previous study^46^ were present in all seven transition lineages close to the transition node and were derived from three HOGs that emerged before the transition node (Fig. 5A). The proportion of all aquaporins that were present differed for each lineage which was expected owing to the diversity of their distribution across eukaryotes^48^. Finally, Hox genes derived from a single HOG were present in all animal and fungal lineages.

A comparable pattern was observed for the independent origins of multicellularity where we examined Beta-catenins, Cadherin protein domains, the Integrin Adhesion Complex^49–51^, NMD machinery and SR splicing factors^52^, as well as lineage-specific regulators such as flocculation genes in yeasts^53^, cyclins in *Chlamydomonas*^54^, and multicellularity-associated genes in fungi^55^ (Table S9). Many of these were found not only in the clades where they had been originally described, but also in others that independently evolved multicellularity. For instance, Dictyostelia and Pezizomycotina gained orthologs of nearly all animal multicellularity-associated genes (except Beta-catenins), while Embryophyta gained all but the Cadherin domains^50^ and the Integrin Adhesion Complex (Fig. 5B; Table S10). Adhesion and signaling systems showed sharp, statistically supported gains near the metazoan transition, whereas cyclins and flocculation-related proteins in plants and fungi increased more gradually, with significant nodes only in a few lineages.

In both phenotypic transitions, we observed that the HOGs containing these orthologs often originated well before the actual phenotypic transition nodes. This pattern suggests that ancient gene families provided a shared genomic background that was independently co-opted for similar adaptive functions. For example, the presence of Hox genes and Aquaporins reflects expansions or lineage-specific deployments of older gene families. Indeed, a recent study showed that functions associated with more than half of all human protein-coding genes have ancient origins, particularly at the root of all eukaryotes^56^. Thus, while individual transitions and species may recruit different gene sets, they frequently draw from a common pool of conserved functions. These analyses show that genes associated with major evolutionary transitions are seldom lineage-specific innovations. Instead, they belong to older gene families that originated well before the transitions themselves and were later co-opted for similar adaptive roles.

### A web application for exploring evolutionary histories

To make our evolutionary inferences widely accessible and to facilitate exploration of both gene- and function-level reconstructions, we developed an interactive web application for browsing evolutionary histories of genes, domains, and pathways generated in this study. This resource (available at https://funcevol.russelllab.org) complements the analyses presented here by enabling customized, exploratory queries across five data types:

1. **Taxon.** For any taxon defined at a node of our species tree (see Materials and Methods), users can view the number and identity of genes, domains, and pathways gained at that node and its descendants, as well as the corresponding losses for genes and domains.
2. **Gene.** For genes from 31 model species, the application displays their presence/absence across all sampled genomes and their inferred states at every ancestral node. Users can choose to trace either the orthologs of a gene or the broader HOG to which it belongs.
3. **Function.** Users can view aggregated histories of functional categories—GO terms, KEGG pathways, and Reactome pathways—facilitated by combining the evolutionary histories of their associated genes.
4. **Keyword.** Free-text queries allow retrieval of summarized evolutionary histories of all genes annotated with a chosen keyword, offering flexible, annotation-driven exploration.
5. **Gene list.** Users can input a custom list of genes (e.g., UniProt accessions) and obtain a summarized evolutionary history of the set, enabling gene-centric analyses that extend beyond curated functional databases.

The web application brings together all reconstructed gene, domain, and pathway histories in a single, queryable platform. Users can navigate from individual genes to higher functional levels, visualize where specific innovations occurred along the phylogeny, and compare patterns across taxa. By linking phylogenomic inferences with functional annotations, the resource provides a reproducible and transparent means to explore the evolutionary signals described here and to generate new hypotheses about gene and function evolution.

## Discussion

Modern genome annotation pipelines combine automated rules and curated knowledge to assign functions to genes^57–61^. Although annotations remain uneven and biased towards model organisms, we hypothesized that they already contain enough information to recover biologically meaningful signals when embedded in a phylogenomic framework. In fact, the availability of accurate and curated annotations for model organisms provided helpful signals at nodes in our phylogenetic tree that were ancestral to those species. As functional genomics and community curation expand, these resources will only become richer and more reliable. Our results show that, even with current biases, annotations can illuminate major evolutionary patterns. Functional enrichments at key nodes closely align with traits historically associated with those clades. For example, developmental processes in metazoans, light and stress responses in land plants, and adhesion and signaling in multicellular fungi. This highlights the untapped potential of existing annotations as predictive tools in comparative and evolutionary genomics.

The central novelty of our study is a top-down framework for evolutionary functional genomics. Instead of tracing one clade or function in isolation, we reconstructed the emergence of all annotated functions across a de novo, genome-derived species tree. This revealed both lineage-specific innovations and convergent enrichment of broad functional categories at major transitions. Terrestrialization in slime molds, plants and animals independently coincided with gains in stress tolerance and developmental genes, while multicellularity in animals, fungi, plants, and slime molds was consistently associated with expansions in cell adhesion and communication functions, even though the underlying gene families differed (e.g. Fig. 4D-E).

Across our analyses of nodes with bursts of gene and function gain and the transitions to land and multicellularity, two nodes warrant special mention: Dictyostelia and Embryophyta where the phenotypic and genomic events coincided. It is either possible that the observed genomic innovations in land plants and slime molds are due to the dual phenotypic transitions that these clades underwent, or the coincidence of these events may also reflect the nature of our species sampling. For instance, a previous study with 208 plant genomes reported two bursts of genomic novelty^39^, which we also observe in two successive plant nodes (Fig. 1), indicating that nuances in either the genomic innovations or the phenotypic transitions (e.g. multiple terrestrial transitions in nematodes^62^ or animals^40^) may be missed due to our relatively sparse sampling. Nevertheless, by sampling broadly rather than deeply, we gained the ability to detect convergence at large evolutionary scales while describing the detailed genomic correlates, including enriched functions and genes gained at each node. Thereby, we complement lineage-focused studies with a broader perspective on shared genomic solutions to similar selective pressures, while acknowledging that lineage-specific novelties, outside the scope of our convergence-focused framework, remain equally important. Together these approaches underscore the value of coupling phylogenomic inference with functional annotation to reveal both convergent and divergent evolutionary strategies.

We also note that gene family ancestral reconstruction inherently reveals both gains and losses; and here we largely focused on gains to examine the emergence of functions, though losses are also detectable (Fig. S3) and can be explored through our web application. Inferences of when genes were gained depend on phylogenetic sampling^63^, and undersampling of certain clades may exaggerate apparent bursts of innovation. Hence, we interpreted all events using functional enrichment and clustering rather than overinterpreting the exact genes and their individual annotations. We also carried out different kinds of robustness analyses that indicate sparser species samplings and other models of ancestral reconstruction do not broadly affect our inferences (Materials and Methods; Fig. S9). Of course, artifacts such as horizontal gene transfer in prokaryotes and uncertainties in deep species relationships remain challenges that future iterations of this framework will address.

Despite these limitations, our analyses demonstrate when the seemingly disparate fields of phylogenetics and functional genomics^37^ are integrated, we can capture both expected and hidden signatures of evolutionary innovation and convergences. Any newly sequenced genome can now be easily placed into this functional phylogenetic framework, revealing both its unique and conserved functional innovations. This provides an immediate launch point for hypothesis generation in functional biology, development, and evolutionary studies of newly sequenced species. Looking forward, integrating standardized phenotypic data with these reconstructions will enable comprehensive genotype–phenotype maps of life, linking genomic changes to traits across deep evolutionary time.

## Materials and Methods

All analyses were done using the R programming language^64^ and the tidyverse set of packages^65^ unless noted otherwise. Most of our large data tables were added to a PostgreSQL database that was queried in R using the dbplyr^66^ and RPostgres^67^ packages. For tree manipulations and visualizations, we used packages such as phytools^68^, ape^69^, ggtree^70^, cowplot^71^, patchwork^72^, and RColorBrewer^73^.

### Proteomes, pathways and functional annotations

We sampled 508 species from across the tree of life (superkingdom Archaea, Bacteria and Eukaryota) from the full list of Uniprot reference proteomes by trying to keep at least one representative from a taxonomic class while proportionately sampling larger clades (proteomes downloaded April 2021, Table S1). We downloaded the canonical protein sequences from each proteome (4,543,007 proteins in total) and used the Uniprot accessions as the anchor point for all subsequent analyses. Next, we downloaded the Uniprot accessions of all protein components for every pathway in KEGG (last updated July 2023) and in Reactome (downloaded July 2022). For each KEGG pathway, we also downloaded the “kgml” graph files associated with every pathway. Using the KEGGgraph R package^74^, we then extracted the components (nodes) connections (edges) in each pathway, be it protein-protein or protein-chemical and all the reactions catalyzed by different enzymes. For GO terms, we extracted all the three GO category terms for nearly every protein in our dataset (4.53 million out of 4.54 million proteins) from the Uniprot knowledgebase file “uniprot_sprot.dat.gz” (August 2021) using the Bio.SwissProt module in Biopython. For each term detected, we also captured the complete lineage using the GO obo file (https://purl.obolibrary.org/obo/go.obo) and the ontologyIndex R package^75^.

### Inferring orthologs and phylogenetic tree building

We used the OrthoFinder (versions 2.5.2 – 2.5.5)^29^ program with the option -og to first infer the Orthogroups (OGs) for all of our proteomes. OrthoFinder has been shown as one of the highest performing orthology programs^76^, it uses a phylogenetic framework and definition of orthology and is scalable for a large number of genomes. Overall, 91% of the 4.54 million proteins in our dataset were assigned to 154k OGs (Table S11). We determined the OGs that had <= 10 gene copies in at least 50% of the species. We built the multiple sequence alignments of each of the fasta sequences of the OGs using MAFFT-linsi^77^ with the option --anysymbol and built gene trees for each one of them using IQ-tree^78^ with the option of model testing with the LG, WAG and JTT models^79^.

Using the gene trees, we removed paralogs using a custom python script; briefly, we checked if the paralogs from a species are monophyletic. If so, we only kept the paralog with the shortest branch length, else, we dropped all paralogs. We then resampled the OG fasta files by keeping one paralog per species and determined the OGs that had a sequence from at least 249 species (roughly 50% of all species). We aligned the 239 selected OG fasta files using MAFFT-linsi, removed positions with large number of gaps using trimAL^80^ with option –gappyout, concatenated the alignments and used FastTree^81,82^ with the options -lg (for the LG model) and -cat 10 (for 10 rate categories) to get the species tree. We chose the LG model and 10 rate categories based on the model testing done for each OG gene tree mentioned above. FastTree uses the Shimodaira-Hasegawa test^83^ to calculate local branch support values by testing all possible alternative topologies and 1000 resamplings of the alignments (Fig. S1). A similar exercise of species tree building was also carried out using the IQ-tree^78^ program, however this yielded a tree with fundamental placement errors such as the nesting of Archaea within Bacteria. This is likely because IQ-tree converged on a local maxima for likelihood (Likelihood = −25290808.895) whereas the tree from FastTree achieved a higher maximum likelihood (Likelihood = −25203470.883), even though it only approximates the maximum likelihood, thus alleviating this issue. Finally, this species tree was used to split orthologs from paralogs, and generate hierarchical orthologous groups (HOGs) by using the - fg, -s, and -y options in OrthoFinder.

### Taxonomy of tree nodes

For every node, we sought to assign taxonomic information and obtained the entire NCBI taxonomy lineage for each species using the taxonomizr R package^84^ (and the “names.dmp” and “nodes.dmp” files from NCBI taxonomy FTP; updated February 2025). We then determined the name from the broadest taxonomic level that encompassed all of the descendant species. In some cases, we also captured names from lower taxonomic levels that encompassed >75% of the descendant species. This approach captured the lowest possible taxonomic identity for all nodes, except where the descendant lineages led to an equal proportion of species from multiple groups (Table S2). In such cases, the node usually had a very broad classification (e.g., superkingdom Eukaryota or Bacteria).

### Assumptions and robustness of methods

We analyzed ∼4.5 million proteins from 508 species using OrthoFinder, which partitions sequences into hierarchical orthologous groups (HOGs) defined at each node of the species tree^29^. Throughout this study, we focused on HOGs inferred at the root of the tree (N0 HOGs).

For functional inferences, we mapped each HOG to curated annotations from Gene Ontology via its gene members. For KEGG and Reactome pathways, the presence of an ortholog of a pathway gene was taken as evidence for that pathway component in a given species. This follows the widely used *ortholog conjecture*, which posits that orthologs are more likely to retain equivalent functions across species than paralogs^85,86^. However, orthologs can also diverge substantially^87^. To avoid over-interpretation, we therefore emphasize aggregate enrichments of functional categories (e.g. GO terms, broader hierarchies of pathways) and the cumulative proportion of pathways gained, rather than claims about the exact roles of individual genes.

Three factors could affect the robustness of our inferences:

1. Proteome quality: We used Uniprot reference proteomes throughout our study to provide a good starting point for annotations. Nevertheless we carried out BUSCO completeness analyses^88^ for 435 of our 508 proteomes (BUSCO failed on the others) using the appropriate lineage datasets. 411 of the 435 proteomes (94.4%) had a completeness score of > 75% with an additional 17 proteomes having a score > 50%. This suggested that the proteomes we used were of high completeness and quality.
2. Inference of gene gains and losses: We used the first reconstructed appearance of a HOG on the species tree as a proxy for gene origination. This provides a conservative estimate and depends on both orthology inference and tree topology. Inferring the ancestral states of HOGs and gene orthologs is a regular task in comparative genomics and a plethora of approaches exist for the same. There are four main approaches that apply to gene families: Wagner Parsimony, Dollo Parsimony, Maximum Likelihood and Phylostratigraphy (i.e., most recent common ancestor of all organisms carrying a gene). Dollo Parsimony has previously been shown to overestimate gene ages^89^ and hence we did not consider this method. We carried out comparative analysis between the other three methods and checked how similar results were. Our method of interest was the Wagner Parsimony with an equal probability of gains and losses (called g1), and we compared this to the Wagner Parsimony with twice as likely to infer gains than losses (called g0.5), with the Maximum Likelihood method (ML) and phylostratigraphy (mrca). When we analyzed the identity of nodes where all of our HOGs were gained using these methods, we observed a strong Jaccard similarity between the g1, g0.5 and ML methods (> 0.9, Fig S9A). The mrca method had very low scores comparatively and this is to be expected as it can often overestimate the gain of a HOG as being on a deeper node without considering multiple gains. Hence, we carried out all analyses in our study using the g1 method of inferring gene gains and losses.
3. Phylogenetic sampling: Inferences of gene gain depend on the taxa sampled. Undersampling of some clades can exaggerate apparent bursts of innovation. Hence, we carried out an analysis by sampling 203 (i.e., 40%), 254 (50%), 305 (60%), 356 (70%), 406 (80%), and 457 (90%) species randomly and checking if this affected our inferences of HOG gains. Across five samplings per run, we observed that the median Jaccard Index was already 1.0 for the sampling with 356 species when we compared the identity of nodes where each HOG was gained in our complete set (Fig. S9B). There was some variance in the Jaccard Index till the sampling with 406 species, but the sampling with 457 species had a complete overlap for the detected HOGs. Of course, the number of HOGs detected varied by sampling and we captured an average of ∼90% of all 166k HOGs in the sampling with 457 species. Overall, this showed that sparser species samplings would not have broadly affected our inferences of HOG gains.

Together, these safeguards ensured that while the precise timing of particular events may be revised with deeper phylogenetic samplings of individual clades, the overall convergence patterns reported at the scale here are robust to current limitations.

### Ancestral reconstruction of genes/pathways/domains

As mentioned above, we assumed that each HOG was a single gene at some point in evolutionary history and duplicated/diverged further^90^. We used the N0.HOGs from OrthoFinder and their presence/absence in every species in our study to carry out ancestral reconstruction. We also created similar presence/absence matrices for the orthologs of every gene calculated above. For KEGG pathways, we used the API to download the KEGG orthology groups associated with all pathways and hierarchies (https://rest.kegg.jp/link/path/ko) and the Uniprot accessions present in every KEGG orthology group. To make a presence/absence matrix for each KEGG orthology group in every pathway and hierarchy, we also extended the KEGG orthology using our own orthologs and then calculated the species where each group was present/absent. We needed to be careful here and not imply the unknown presence of a KEGG pathway in some organisms as a result of using our orthology. Hence, we forced the KEGG orthology groups associated with a pathway to be absent in an organism if the pathway also did not exist for that organism in KEGG. In the case of Pfam domains, we determined the Pfam domain content for all Uniprot accessions in our dataset and calculated the presence/absence of each domain in our species.

As mentioned before, for ancestral reconstruction in each case, we used the Count program^91^ with the Wagner parsimony^92^ option and gain/loss penalty equal to 1 meaning both are equally possible. From the output that the Count program provided, we extracted the states of each entity at every node in the tree and noted the descendant nodes where the state changed from absent (in the parent node) to present (in the descendant node) as “gain nodes” and “loss nodes” for the opposite.

### Biological entity counts

We counted the number of genes and Pfam domains, gained at every node in the tree by checking if the state of the parent node was “absent” (0) and the node of interest was “present” (1). For KEGG/Reactome pathways, we determined the proportion of components “present” at every node of the tree and used the difference between the proportion at a node and the proportion at the parent node as the proportion gained at each node (Table S1, Table S2).

### Ancestral reconstruction of phenotypes

For phenotypes, we decided to focus only on eukaryotes as the encoding of phenotypes was distinct from bacteria. While bacterial habitats could be inferred for most species in our study, their habitats are very dynamic and non-discrete in many cases. Conversely, in eukaryotes we could use a broad classification. For instance, we considered only the primary habitat of the adult where it spends the most time in nature. Only frogs and the duck-billed platypus were considered amphibians. All eukaryotes that were pathogens or commensals were called “Host-associated”. The terrestrial label also included airborne organisms. The cellularity phenotype was relatively straightforward to encode (Table S1).

Using the trait matrix for the tips of the eukaryotic subtree in our study (199 tips), we carried out stochastic character mapping^93^ using the “make.simmap” function in the “phytools” package of R. In each case we used the Bayesian “mcmc” option to calculate the transition matrix with the equal rates (ER) model and the number of simulations being 100. We forced the eukaryotic root to be aquatic and unicellular and allowed the function to estimate the probabilities of each trait category at every node of the tree. To assign an ancestral state to the nodes in the tree, we used a posterior probability cutoff of 0.5 for habitat data and 0.75 for the cellularity data i.e. if any category had a probability >= the cutoff, we assigned that as the state. Otherwise, the state of the parent node was transferred to the descendant node. We then determined the nodes that had at least three descendant species where the states switched from aquatic to terrestrial or unicellular to multicellular.

### GO term enrichment and clustering

Throughout the manuscript, we have carried out enrichment of GO terms using a modified Fisher’s test. For any node of interest, we made a 2×2 contingency table as follows:

|  | At Node | Not at Node |
| --- | --- | --- |
| With GO term | # unique HOGs | # unique HOGs |
| Without GO term | # unique HOGs | # unique HOGs |

By considering the number of unique HOGs to represent genes that contain a term, we were able to ensure independence for the gene numbers since each HOG can be thought of as an independent entity. We believe this approach was extremely powerful and helped capture the results in our study. The p-values were corrected using the Benjamini-Hochberg method and enrichment was filtered by keeping GO terms with an Odds Ratio > 1 and an adjusted p-value < 0.05. To get the top terms in Fig. 2, we had to consider the hierarchical structure of the GO term data and hence we removed some descendant terms in the filtered significant lists by keeping only the parent terms for descendant terms seen in the list.

For the GO term clustering in Fig. 3, we analyzed each GO category (BP, MF, CC) separately. We focused on the enriched terms at each of the transition nodes which were determined by taking the terms having p-value < 0.05, a finite odds ratio > 1 and at least five HOGs with the term. We used the “term_sim” function from the simona package^94^ in R with the “Sim_Lin_1998” method^38^. We chose the Lin method since it calculated semantic similarity considering the hierarchical structure of GO and by scoring pairs of terms with the most informative common parent term highly. We clustered the terms using the “cluster_terms” function from the simplifyEnrichment package^95^ with the “louvain” method and a resolution parameter of 1. We determined the top five keywords in each cluster using the “keyword_enrichment_from_GO” function from the simplifyEnrichment package with default parameters. This function carries out a Fisher’s test for the words in a GO term list and arranges them by adjusted p-values. Lastly, we used the full lineage of each term in a cluster to determine an informative cluster name (Table S3, Table S5). Within each cluster, we calculated the mean of the Log2 Odds Ratio for the terms at the transition node and compared them with the mean of the Log2 Odds ratio of all the parent nodes per lineage. Here, we used a t-test with an alternate hypothesis that the ratio at the transition node was greater than the parent nodes.

### Term of interest searches

We exploited the hierarchical nature of GO terms to enhance the searching of a function of interest. Each keyword was used as a regular expression to search a SQL table containing the entire hierarchy of all terms associated with all genes in our study. As a result, even genes that may have the function of interest as a broader parent term can be included to understand the evolutionary history of the keyword i.e. function itself. This is especially useful when a specific GO term assigned to a gene may not necessarily contain the keyword of interest in the term name. For all the genes captured for a search, we took the nodes where they were gained and to reduce the noise in the taxonomic assignments, assigned nodes with taxonomy levels lower than order up to the order level and combined the calculations for each taxonomic name. This approach was in general conservative and sometimes captured minute presences of GO terms in spurious taxa (Fig. S10). To circumvent these, we used coarse intervals for the % genes gained at a node (2%) and also calculated the z-scores of the % gained for a taxon in relation to the mean for a search term and determined the taxa which had a z-score > 1.96 (asterisks in Fig. 2E).

### Transition node orthologs

We sourced the known genes in the phenotypic transitions and made some choices that are detailed in Tables S7 and S9. For the unicellular to multicellular transition, there were other known genes in the animal ancestor that have been attributed to be involved in the unicellular to multicellular transition such as Wnt, TGF-ß, SMADs, Frizzled, dishevelled, Hemicentins, Fermitins and Dystroglycans, however these exclusively emerged in the Metazoa lineage in our analysis and were not shown in the main figure (Fig. S8). In a few cases, we searched the gene descriptions in our database of genes to get a set of starting genes.

In each case, we checked the gene/HOG/protein domain status at the nodes in each transition lineage from the ancestral reconstruction output described above. For the gene status at the tips, we also considered only the genes from species that were terrestrial or multicellular as applicable. The % of orthologs present at each node/tip was calculated as the sum of genes in a set that were present divided by the number of genes.

We converted our tree into a cladogram i.e. ignored branch lengths so as to make all the tips equidistant from the root of the tree (using the “chronos” function in the phytools package). This allowed us to scale the distance of each node/tip between 0 and 1 where 0 represents the root and 1 represents any tips. We also forced all transition nodes to be at 0.5 for better visual representation and analysis. For the % orthologs present values along a lineage, we fitted four linear models for scaled distance intervals of [0-0.25], (0.25-0.5], (0.5-0.75] and (0.75-1]. We took the fitted residuals to draw the lines in Fig. 3A and 3B.

## Acknowledgements

We would like to thank Arnau Sebé-Pedrós, Jan Korbel, Supriya Khedkar and the Russell group for feedback and suggestions. We thank Damien DeVienne and David Emms for help with the LifeMap and OrthoFinder tools. We thank Denise Duma for introducing semantic similarity. We acknowledge the use of chatGPT for checking grammar, overall clarity and flow of the manuscript text.

## Funding

Wellcome Trust UK 210585/A/18/Z (GDD)

German Federal Ministry of Research, Technology and Space (BMBF) PreDiCT, FKZ-01GM2306 (GDD)

European Union Horizon 2020 PrecisionTox project, Grant Agreement Number 965405 (JCGS, MEC)

Swedish Research Council Grant VR2018-05882, UmU project number 322045306 (JCGS)

## Author contributions

Conceptualization: RBR, GDD, MJT, RG, JKC

Methodology: GDD, RBR, PN, MJT

Data Curation: GDD, JCGS, MEC, PN, RBR Software: GDD, JCGS, PN

Investigation: GDD, RBR Visualization: GDD, RBR Funding acquisition: RBR Project administration: RBR Supervision: RBR

Writing – original draft: GDD, RBR

Writing – review & editing: GDD, JCGS, MJT, RG, JKC, RBR

## Competing interests

Authors declare that they have no competing interests.

## Data and materials availability

All data are available in the main text or the supplementary materials, an online web application and scripts via a GitHub repository (https://github.com/gauravdiwan89/2025_Diwan_etal_FuncEvol).

## Supplementary Materials

Figs. S1 to S10

Tables S1 to S11 (as one file)

## Notes

https://funcevol.russelllab.org/

### Competing Interest Statement

The authors have declared no competing interest.

### Summary of Updates

Updated manuscript based on peer reviews. Added a new figure and split an earlier figure to improve the narrative. Many more explanations in the main text and more methodological details provided.

