## Supplementary File for "Phylogeny and gene function integration uncovers multiple convergences in multicellular and terrestrial transitions"

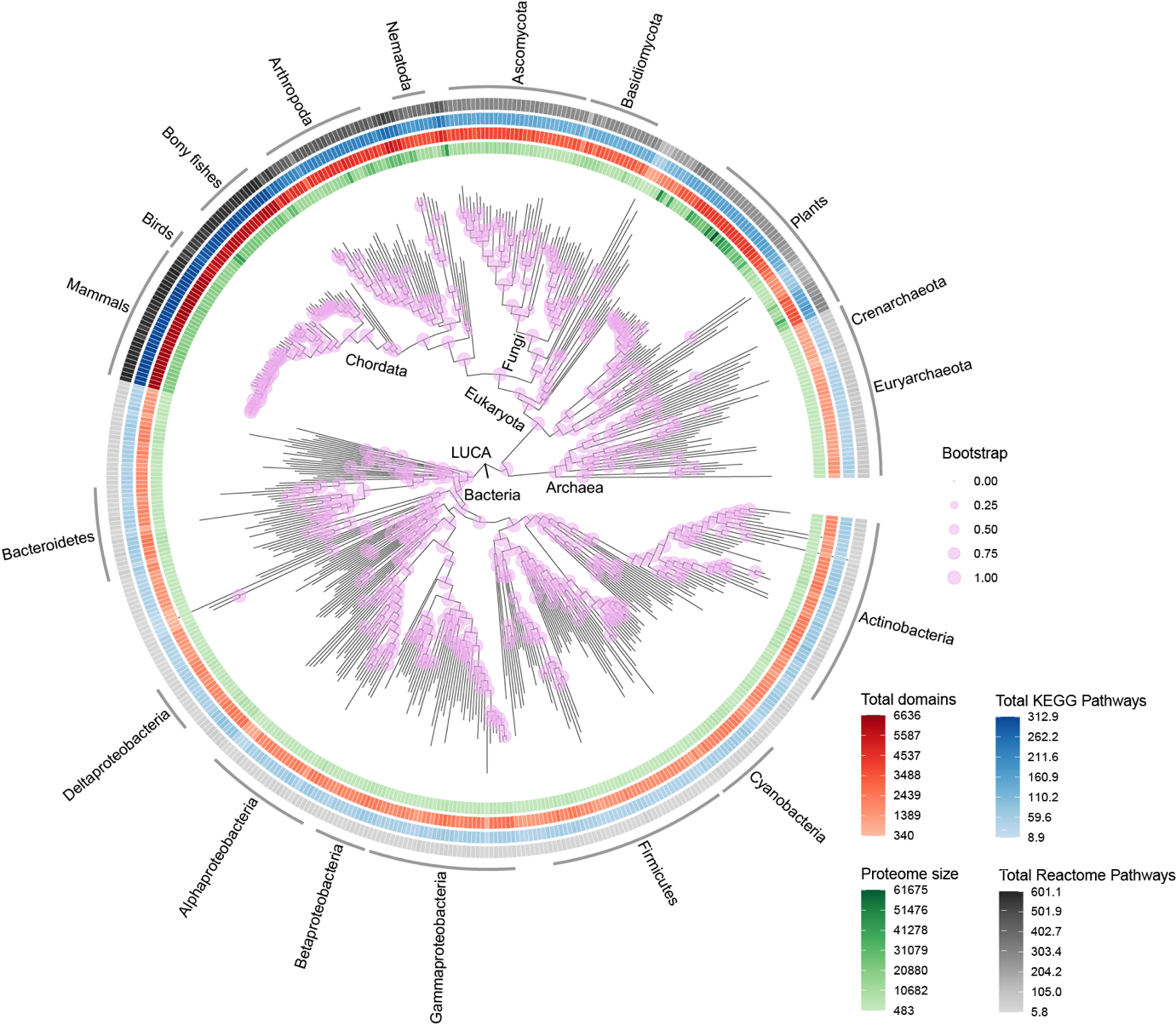

**Fig. S1:** Phylogenetic tree showing the relationships between 508 species from across the tree of life. The nodes have been annotated with points where the size is proportional to the bootstrap values (purple). The colored rings around the tree indicate the number of proteins (shades of green), number of unique Pfam domains (shades of red), total sum of the proportion of all KEGG pathways (shades of blue) and the total sum of the proportion of broad level Reactome pathways (shades of gray).

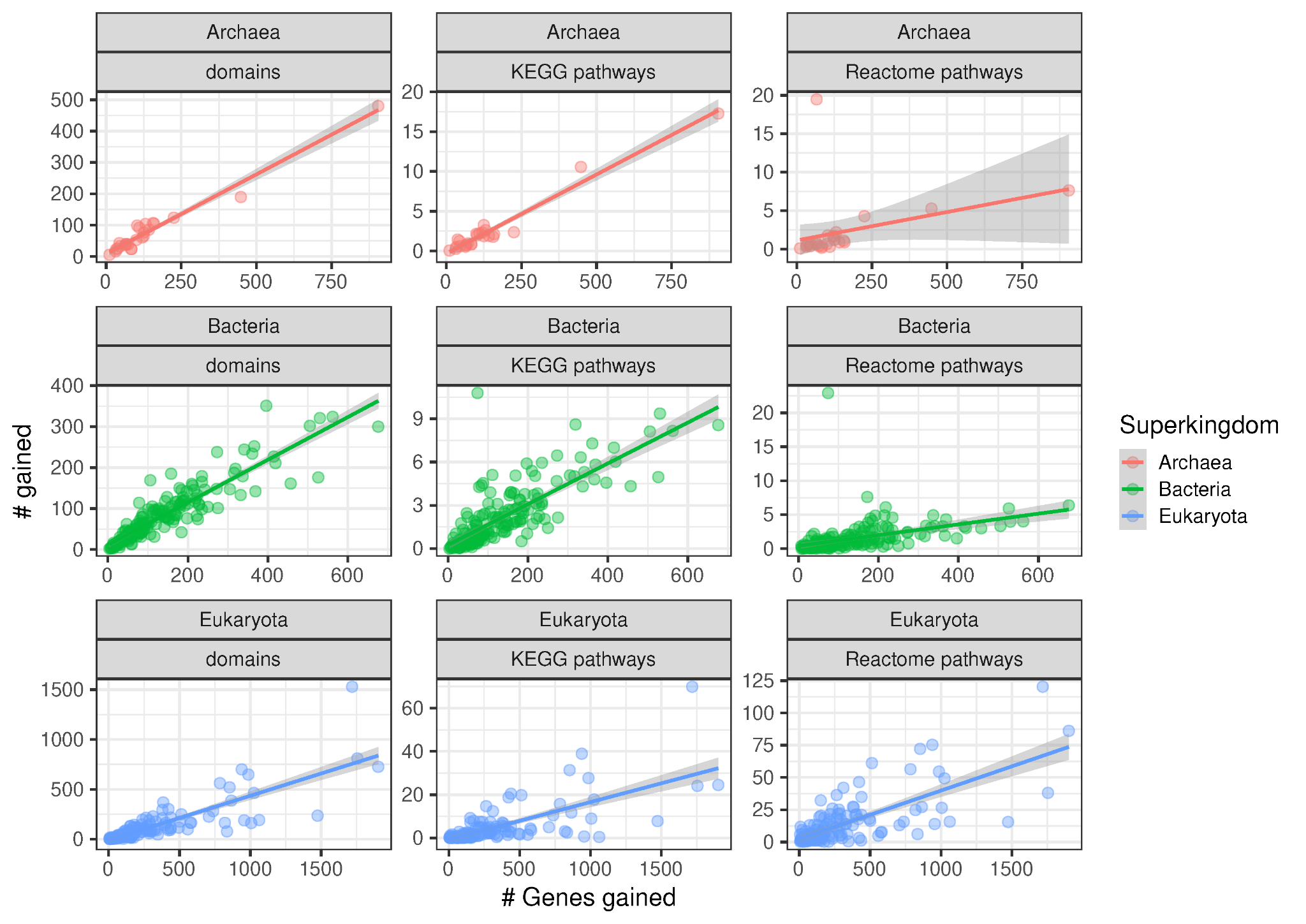

**Fig. S2:** Number of gained Pfam domains, KEGG pathways and Reactome pathways versus number of HOGs gained at every node in the phylogenetic species tree (nodes with >2 descendant species). Linear models were fitted to each category across superkingdoms and the grey regions indicate the standard error of the fit.

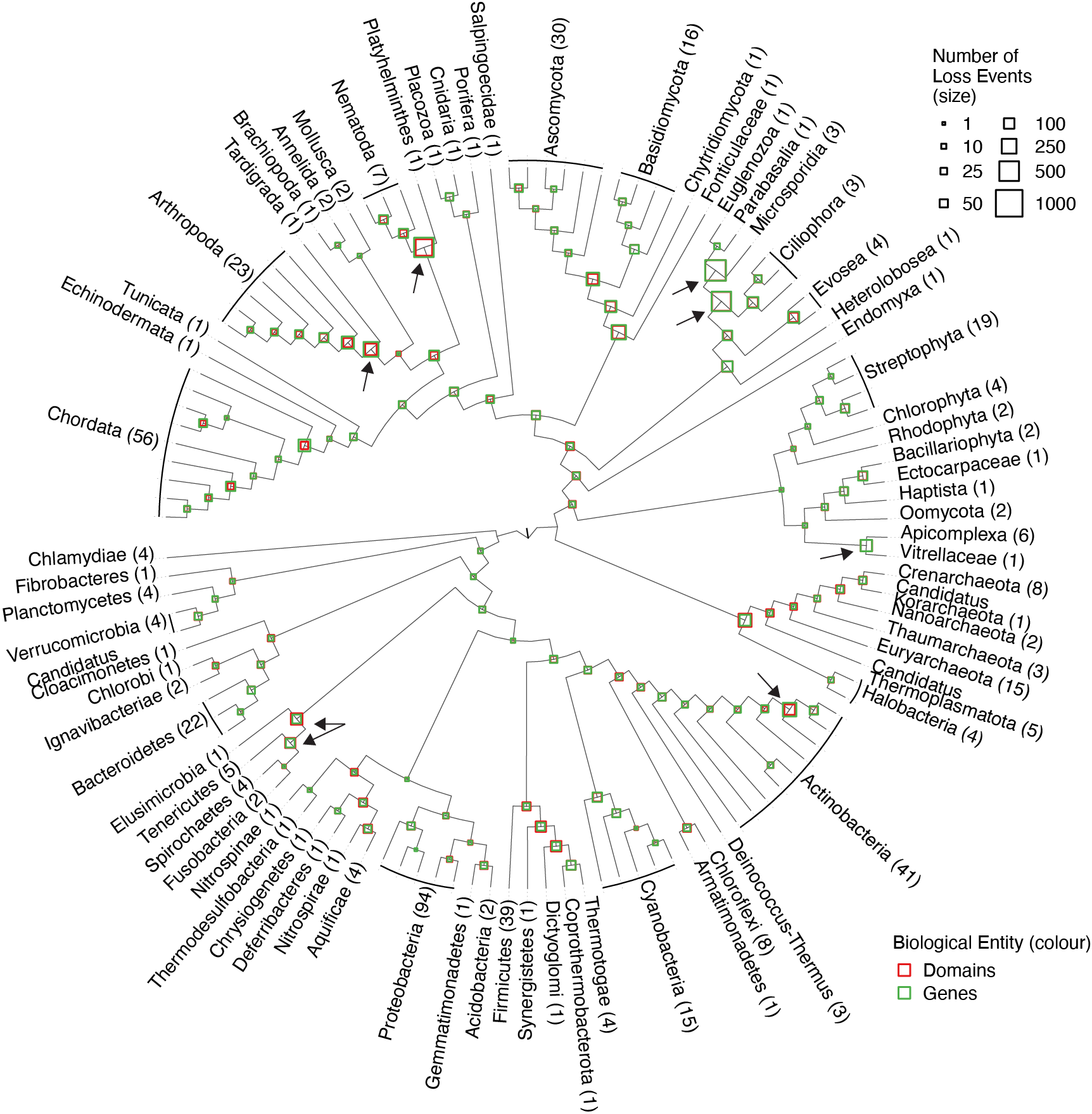

**Fig. S3:** Cladogram similar to Fig. 1 showing the number of HOGs (i.e., genes) and Pfam domains lost at each node where the area of the square points is proportional to the number of losses. Nodes with the largest number of losses have been marked with an arrow and have been described in the main text and in Fig S4B.

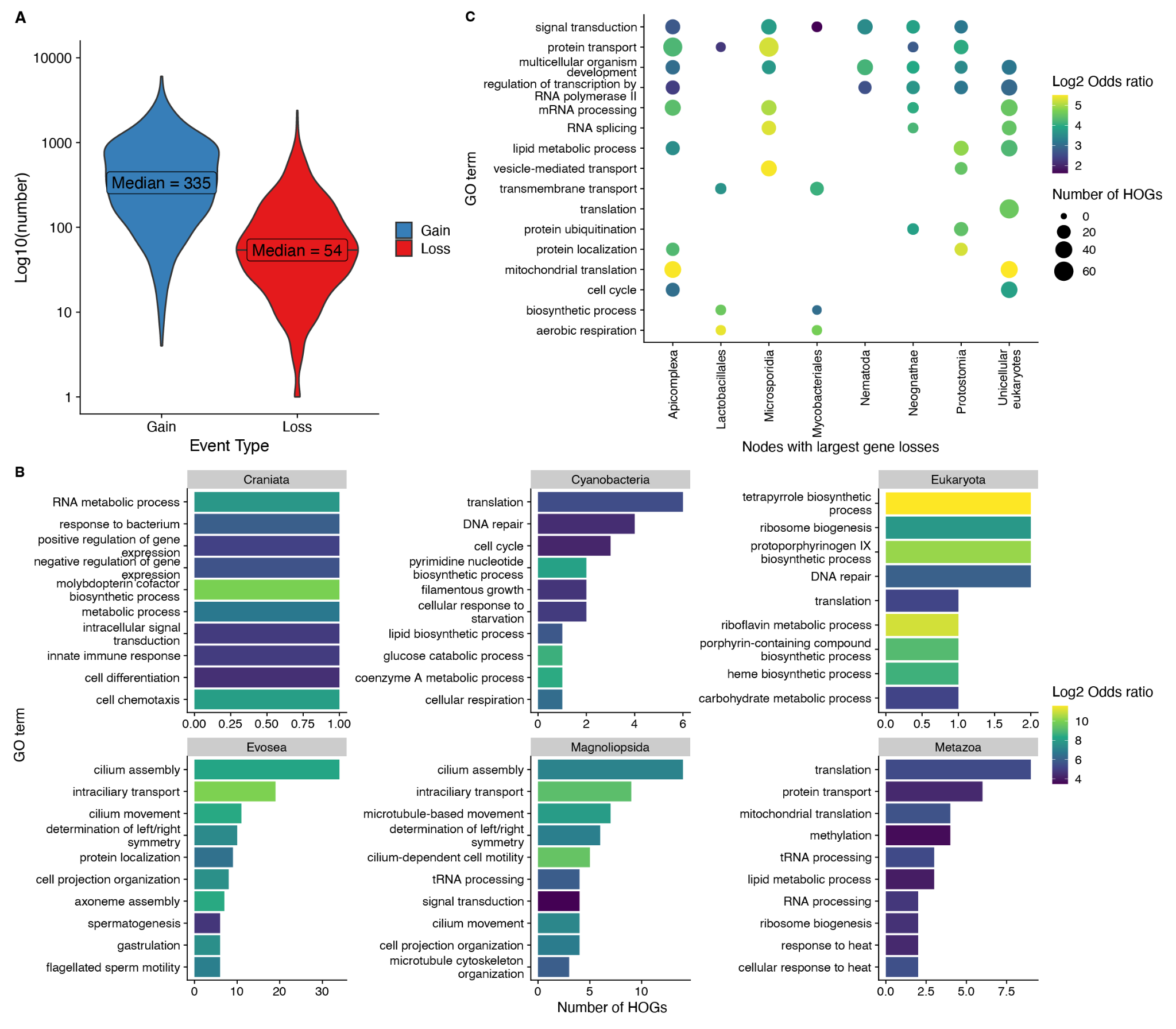

**Fig. S4:A)** Violin plots comparing the number of gains and losses at every node and tip of our complete species tree. The median for each event type is also indicated. **B)** Bar plots showing the number of HOGs with the top ten enriched GO terms at each of the nodes from Fig. 1 and Fig. 2 that had the largest number of gains. The bars have been coloured with the Log Odds Ratio of the enrichment. **C)** GO terms enriched and shared across multiple nodes that showed the largest number of losses (also highlighted in Fig. S3). The size of the points is proportional to the number of HOGs that had the respective term and the colour indicates the Log Odds Ratio of enrichment.

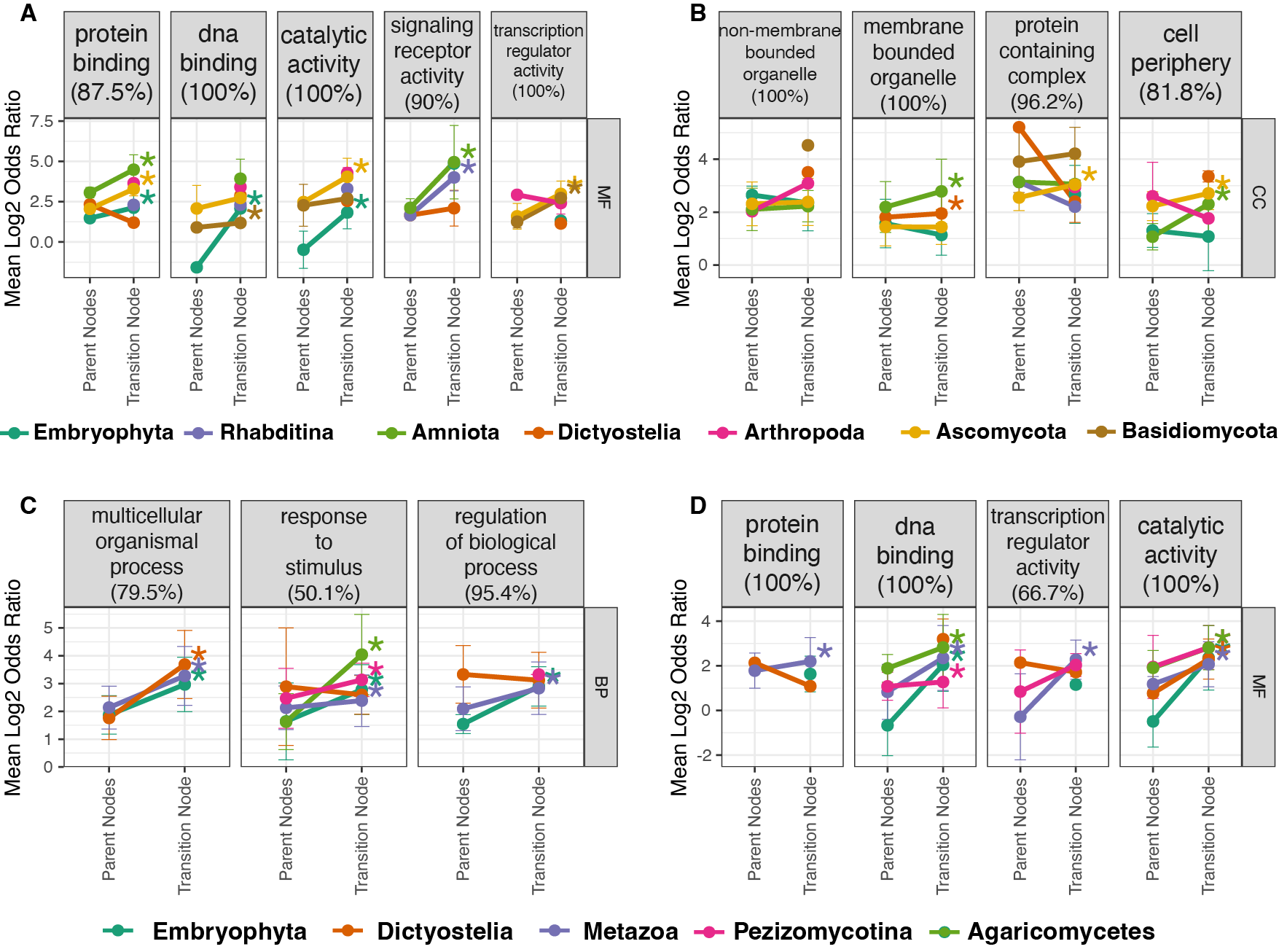

**Fig. S5:** Figures similar to Fig. 4B and 4C showing the enrichment levels of clustered GO terms in the terrestrial transition (panel A and B) and the multicellular transition (panel C and D). These are for terms in the GO categories not represented in the main figure i.e. MF and CC for the terrestrial transition; BP and MF for the multicellular transition. Asterisks indicate significantly larger Log2 Odds Ratios at the transition node compared with the parent nodes in each transition lineage.

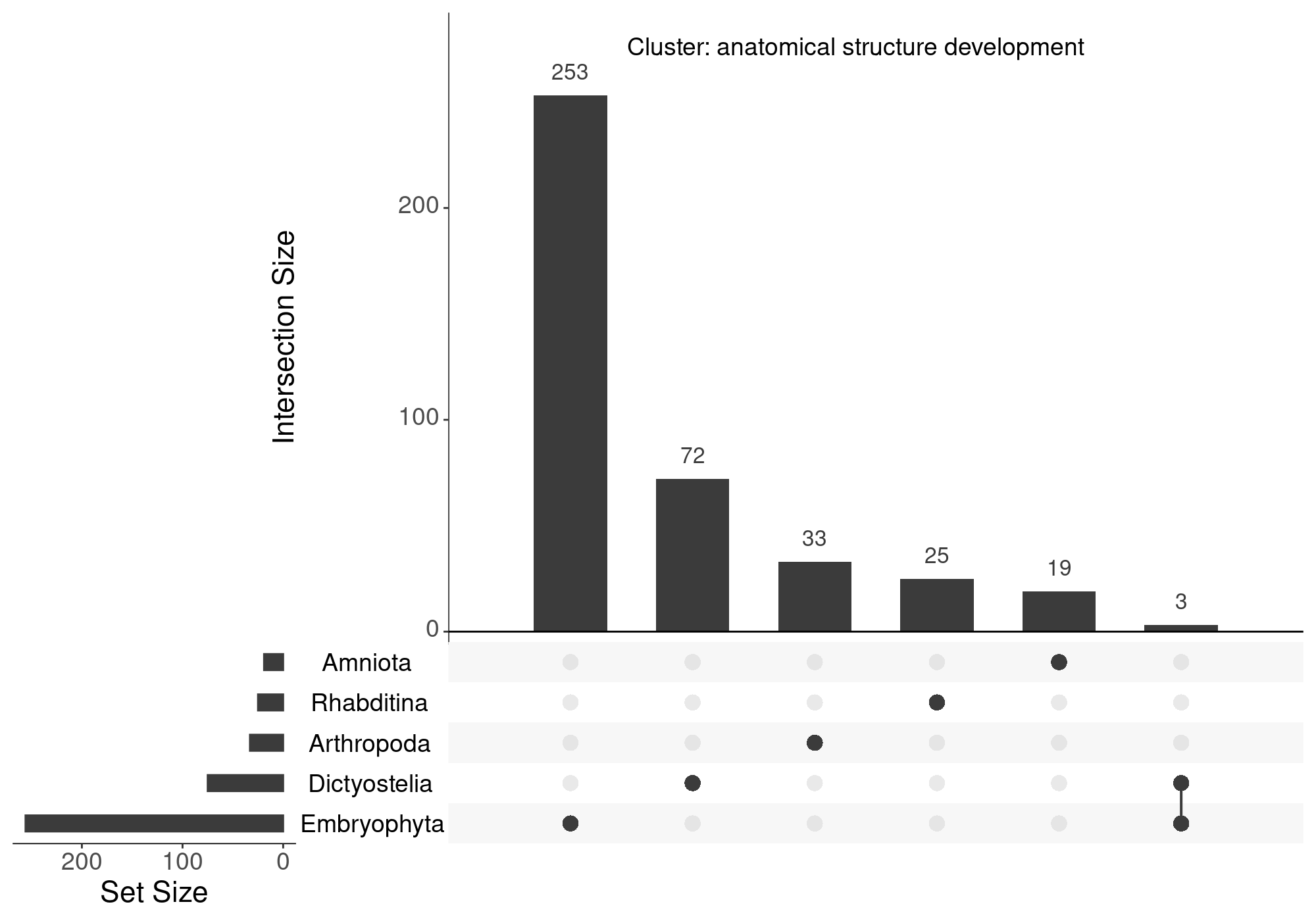

**Fig. S6:** Upset plot showing the number of HOGs emerging at each of the terrestrial transition nodes and if any are shared between the nodes. The HOGs were taken for the GO cluster pertaining to anatomical structure development and showed only a small overlap of three HOGs across Dictyostelia and Embryophyta and no other overlap indicating the independent gain of genes involved in the anatomical structures for each clade. GO clustering enables these genes to be analyzed together.

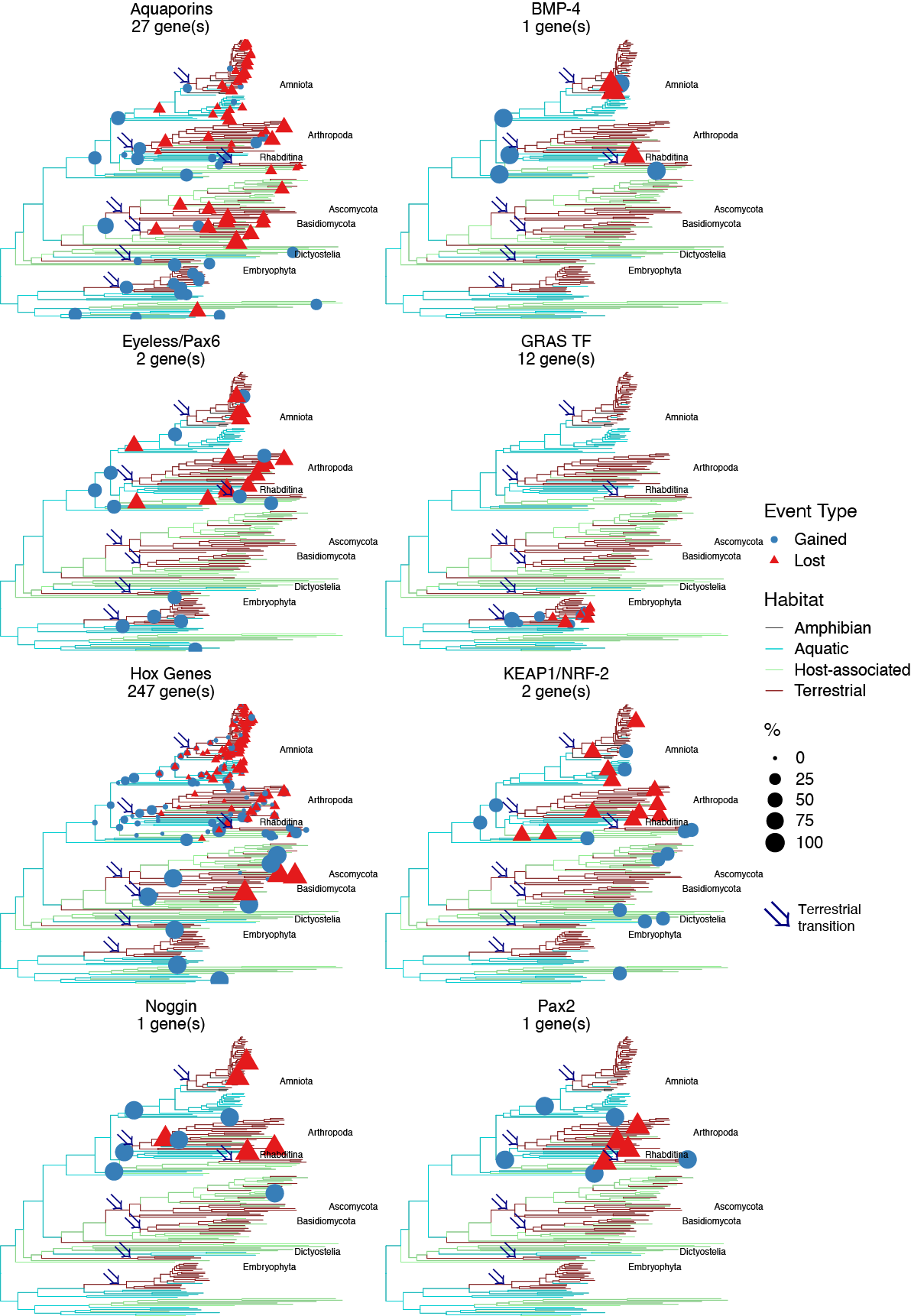

**Fig. S7:** Phylogenetic tree of the eukaryotes in our study with branches coloured by the inferred habitat of the parental nodes. The seven terrestrial transitions are marked with an arrow. Events depicting the gains or losses of ***orthologs*** for known genes (Fig. 3A) are marked by scaled blue filled circles or red filled triangles respectively.

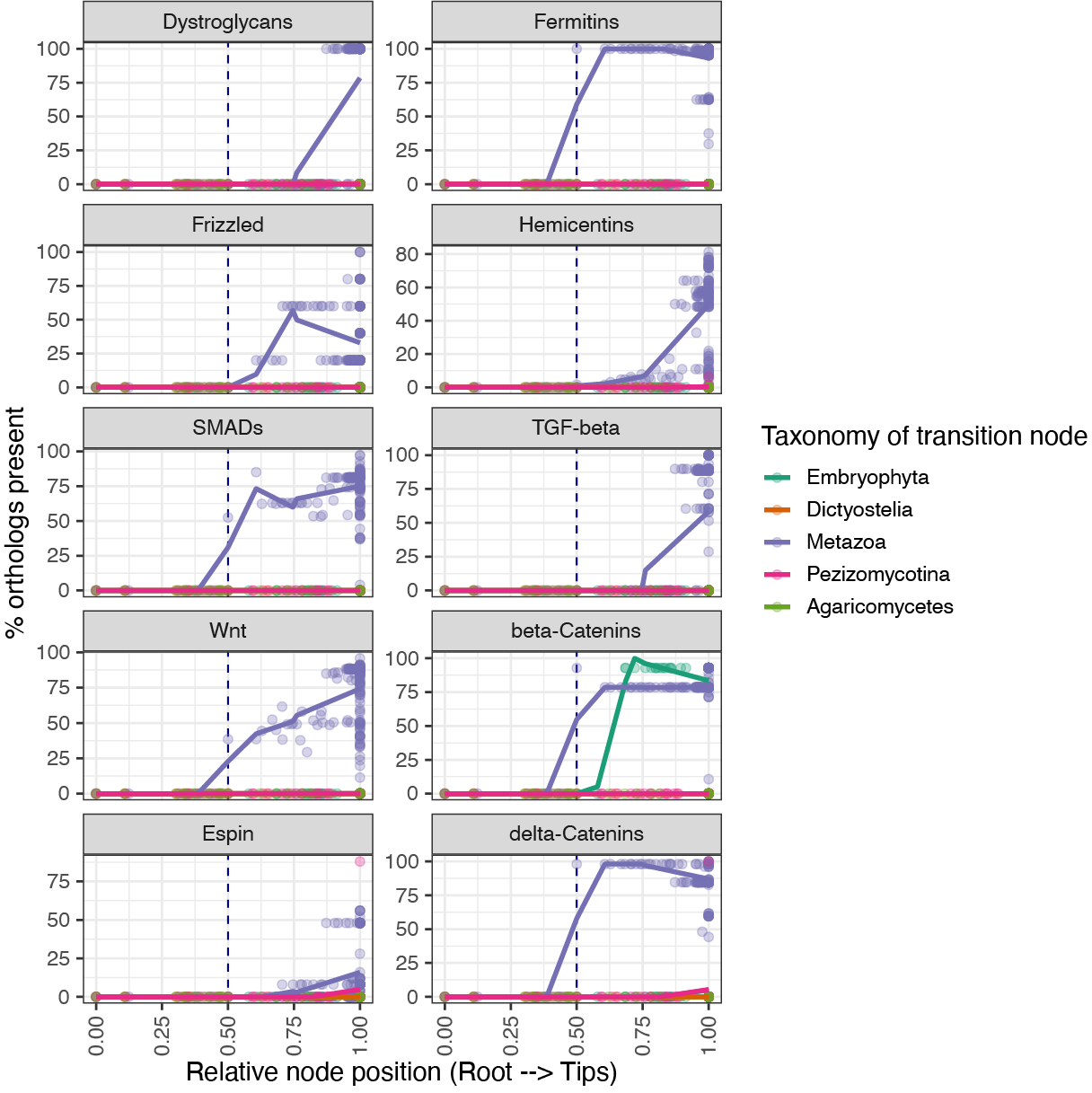

**Fig. S8:** Plot similar to Fig. 5B showing the % of orthologs for genes thought to be present in the animal ancestor at each node of a transition lineage. The distance to the root of each node in a lineage was scaled between 0 and 1 on the x-axis, while the transition node was forced to be at 0.5. Multiple linear models were fitted to the underlying points for each lineage.

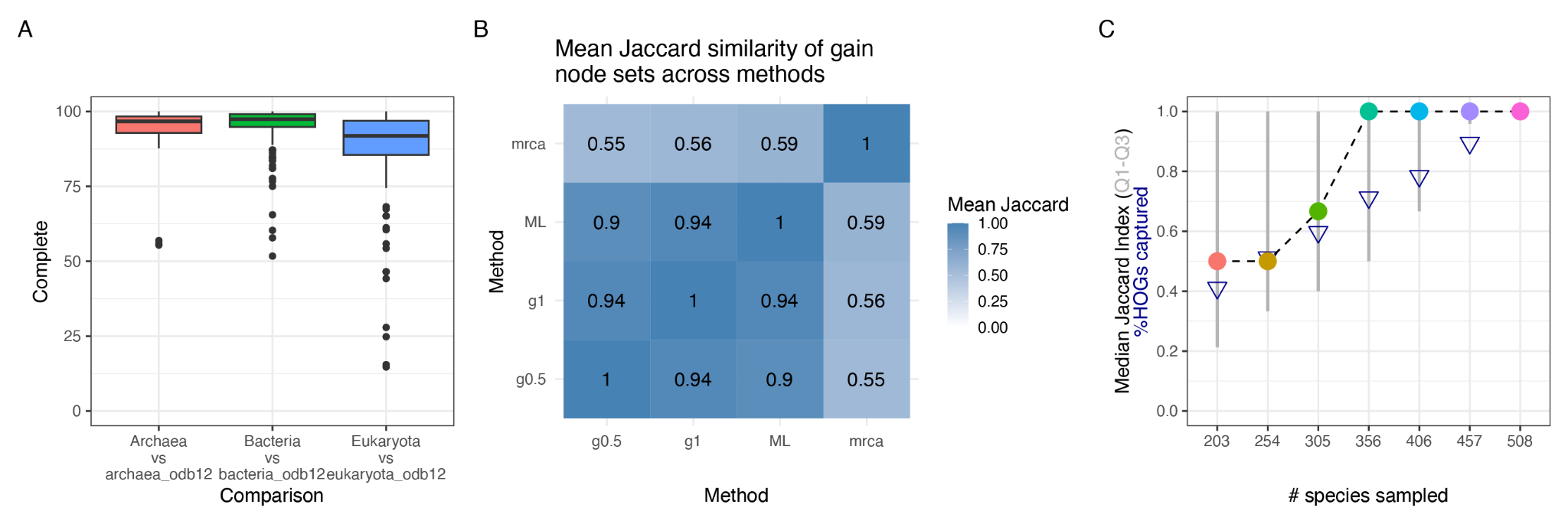

**Fig. S9: A)** Boxplots showing the BUSCO completeness scores for 435 (out of 508) proteomes in our study. The bacterial, eukaryotic and archaeal proteomes were tested against their respective lineage sets. **B)** Heatmap showing the mean jaccard similarities of the gain nodes identified for each HOG across ancestral reconstruction methods. g0.5 = Wagner Parsimony with a gain:loss penalty of 2:1; g1 = Wagner Parsimony with a gain:loss penalty of 1:1; ML = Maximum Likelihood; mrca = Most Recent Common Ancestor of all tips where the HOG was present i.e. a phylostratigraphic inference. **B)** Median Jaccard Index (and Q1 to Q3 quartile values) of the gain nodes identified for each HOG across sparser samplings of the species in our study (done five times for each sampling). Blue triangles indicate the proportion of ~166k HOGs (from 508 species) that were captured in the samplings.

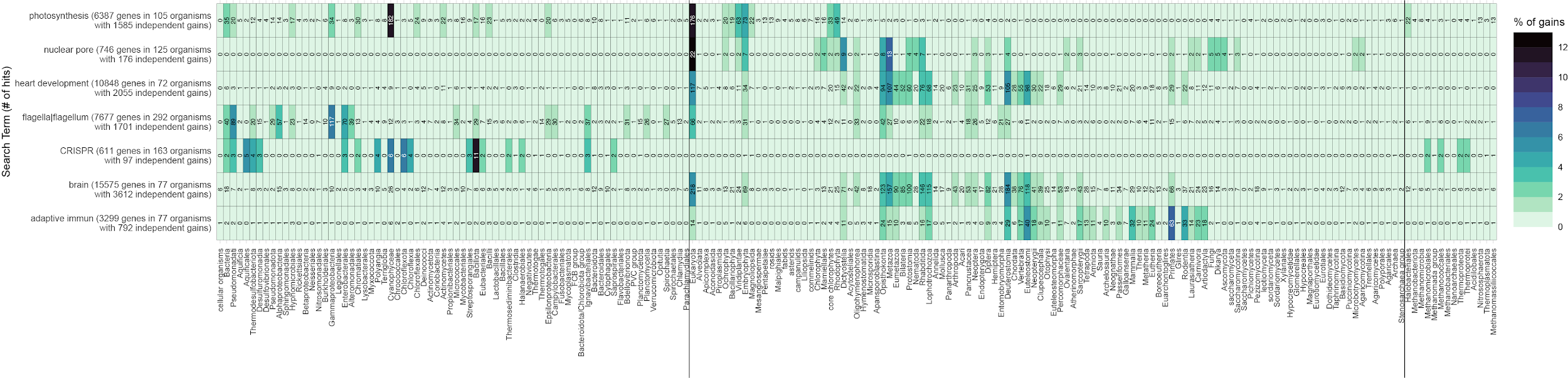

**Fig. S10:** A detailed heatmap, similar to Fig. 2E, showing the % of gains for the genes captured by each search term occurring at each taxon on the x-axis. The numbers in each cell indicate the number of gain events occurring at that node. There is no filtering of taxa in this heatmap and shows the small amount of spurious gain events being inferred in unexpected clades (discussed in Materials and Methods).
